# Signatures of cell death and proliferation in perturbation transcriptomics data - from confounding factor to effective prediction

**DOI:** 10.1101/454348

**Authors:** Bence Szalai, Vigneshwari Subramanian, Róbert Alföldi, László G. Puskás, Julio Saez-Rodriguez

## Abstract

Transcriptomics perturbation signatures are valuable data sources for functional genomic studies. They can be effectively used to identify mechanism of action for new compounds and to infer functional activity of different cellular processes. Linking perturbation signatures to phenotypic studies opens up the possibility to model selected cellular phenotypes from gene expression data and to predict drugs interfering with the phenotype. At the same time, close association of transcriptomics changes with phenotypes can potentially mask the compound specific signatures. By linking perturbation transcriptomics data from the LINCS-L1000 project with cell viability phenotypic information upon genetic (from Achilles project) and chemical (from CTRP screen) perturbations for more than 90,000 signature - cell viability pairs, we show here that a cell death signature is a major factor behind perturbation signatures. We use this relationship to effectively predict cell viability from transcriptomics signatures, and identify and experimentally validate compounds that induce either cell death or proliferation. We also show that cellular toxicity can lead to an unexpected similarity of toxic compound signatures confounding the mechanism of action discovery. Consensus compound signatures predict cell-specific anti-cancer drug sensitivity, even if the drug signature is not measured in the same cell line. These signatures outperform conventional drug-specific features like nominal target and chemical fingerprints. Our results can help removing confounding factors of large scale transcriptomics perturbation screens and show that expression signatures boost prediction of drug sensitivity.

## 1. Introduction

Predicting cellular phenotypes (e.g.: disease states, cancer drug sensitivity etc.) from different high-coverage molecular (‘omics’) data is a key question of current systems biology research. Due to the affordability of its acquisition and the well-established methodologies for analysis, transcriptomics (microarrays or more recently RNA-Seq) is one of the key data sources for these studies (McGettigan, 2013). While basal gene expression can give valuable information about cell state and function, perturbation transcriptomics signatures (i.e., measured gene expression changes after different perturbations such as drugs, gene overexpression or knockdown / knockout) provide additional possibilities to infer cellular function (Lamb et al., 2006). Compounds with similar mechanism of action (MoA) tend to have to similar transcriptomics changes, making perturbation signatures a valuable tool to identify MoA of unknown compounds (Iorio et al., 2010; Lamb et al., 2006; Subramanian et al., 2017). Furthermore, perturbation of different cellular pathways with pathway specific perturbagenes allows the identification of pathway-regulated genes, from which pathway activity can be effectively inferred (Parikh et al., 2010; Schubert et al., 2018).

Small scale studies (comprehensively collected in (Wang et al., 2016)), and the original Connectivity Map study (Lamb et al., 2006) provide rich perturbation signature data. The recent release of the LINCS-L1000 dataset (Subramanian et al., 2017), with more than 1,000,000 signatures, increases these numbers by an order of magnitude. In the LINCS-L1000 screen more than 20,000 different perturbagenes (compounds, shRNAs, etc.) were used in dozens of different cell lines, with different concentrations and perturbation times. Importantly, these high-throughput measurements were possible based on the inexpensive L1000 methodology, that measures only ~1000 (landmark) genes, while the rest of gene expression values were inferred. While this dataset alone opens myriads of possible applications, linking these perturbation signatures with other large scale, phenotypic studies enables to model cellular function on a previously unavailable scale.

Arguably the simplest, but at the same time one of the most important cellular phenotypes, is cell viability - cell death or proliferation. Analysing cell viability data together with perturbation signatures is especially important in cancer research, as it can help to understand the mechanism of anti-cancer drugs, and open new therapeutic possibilities. A recent study (Niepel et al., 2017) analysed about 600 pairs of anti-cancer drugs and breast cancer cell lines where perturbation transcriptomics signatures (using the L1000 methodology) and cell viability were parallely measured, leading to important information regarding cell line specific drug effects and drug synergy. There are also rich sources of other cell viability datasets available - but without the corresponding expression measurements upon perturbation. In particular, preclinical studies like GDSC (Iorio et al., 2016), CTRP (Seashore-Ludlow et al., 2015) or NCI60 (Shoemaker, 2006) generated large scale cell viability datasets following drug (compound) perturbations, to identify potential drug sensitivity biomarkers. Other approaches, like project Achilles (Tsherniak et al., 2017), used shRNA screens and created large scale gene essentiality data sources, with the aim of identifying potential new anti-cancer targets. These datasets partially overlap with the LINCS-L1000, allowing an integrated analysis of perturbation signatures and cell phenotype (Jung et al., 2018).

Another important aspect of cell viability and perturbation signatures is related to the fact that cell death can lead to transcriptomic changes unrelated to the perturbation, and this phenomenon can be a confounding factor to infer mechanism of action. Also one of the original LINCS-L1000 papers (Smith et al., 2017) found that some cell line and perturbation independent factor is responsible for the largest part of variability in the L1000 signatures. This factor has been hypothesised to be related to some general cell biological effect like cell viability or proliferation, but this has not been analysed and thus remains uncertain if this is the case.

In this study we analysed the associations between perturbation signatures and cell viability by matching (same cell line and perturbation) more than 90,000 data points between LINCS-L1000 project (perturbation signature) and the CTRP drug and Achilles shRNA screens (cell viability) - creating the, to our knowledge, largest integrative analysis of gene expression signatures and cell viability. We identified a common “cell death signature” and were able to predict cell viability effectively even across studies from different sources and types of perturbations. By analysing pairwise signature similarities, we found that the “cell death signature” can lead to unexpected similarity between signatures of toxic compounds and thereby can influence the mechanism of action identification. However, using a reduced signature (removing genes showing high correlation with cell viability) we were able to reduce this effect. Our models allowed us to predict cell viability for all the compounds used in the LINCS-L1000 dataset, identifying several potential drugs with death-inducing or pro-growth properties. By using consensus compound signatures and machine learning models, we were able to predict anti-cancer drug sensitivity even in cell lines where the drug signature was not measured, outperforming conventional drug specific features (like nominal drug targets or chemical fingerprints).

## 2. Results

### 2.1 Signatures of cell death in the LINCS-L1000 dataset

To analyse the possible effect of cell death on the perturbation signatures from LINCS-L1000 dataset, we matched instances from LINCS-L1000 (Subramanian et al., 2017) with cell viability data from the Cancer Therapeutics Response Portal (CTRP) (Seashore-Ludlow et al., 2015) and shRNA abundance data from project Achilles (Tsherniak et al., 2017). The matching was done based on cell line, perturbation and, in case of compounds, concentration (Fig. 1A, Methods). While cell viability and shRNA abundance are different metrics, they are related (as both of them are proportional to the number of surviving cells after drug or shRNA treatment), so for simplicity we will refer both as cell viability from now on. For perturbation signatures we included only the actual measured (landmark) genes in this whole study. While in the CTRP and Achilles screen all cell viability values were measured at one time point (72 hours and 40 days / 16 population doublings, respectively) after perturbation, in the LINCS-L1000 dataset perturbation signatures were measured at different time points. Hence, it is possible to match two different LINCS-L1000 signatures (same compound/shRNA and cell line, but different time points) with the same cell viability value (Fig. 1A). Using our matching criteria we were able to compose two datasets: CTRP-L1000 of 16390 matched perturbation signature - cell viability pairs (326 compounds, 48 cell lines, STable 1) and Achilles-L1000 of 77230 matched perturbation signature - cell viability pairs (12925 shRNAs, 11 cell lines, STable 1), resulting in the - to our knowledge - largest current matched perturbation signature - cell viability dataset.

**Figure 1 -.**
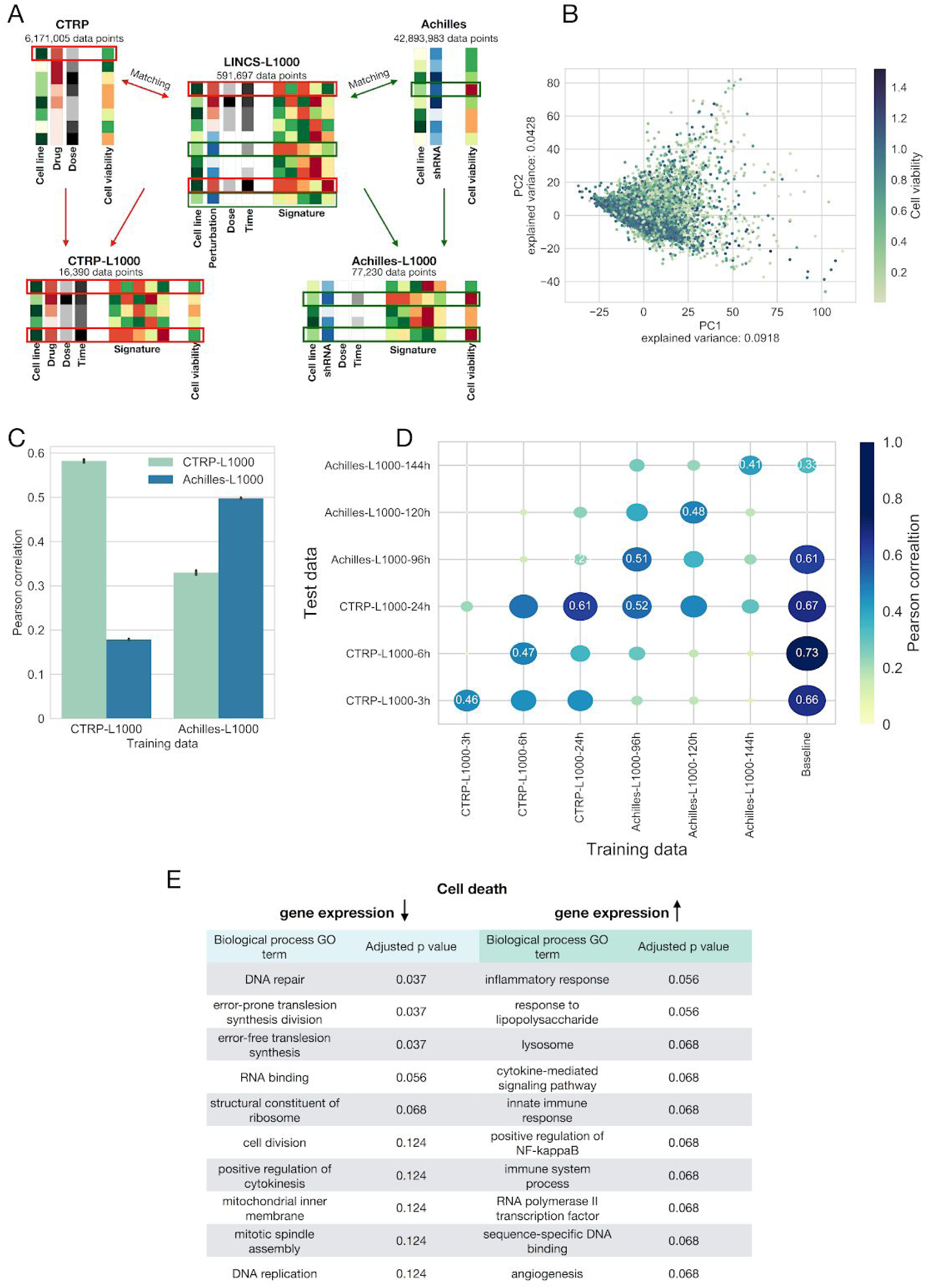
LINCS-L1000 perturbation signatures allow efficient prediction of cell viability. (A) Schematic representation of database matching pipeline. Perturbation signatures from LINCS-L1000 dataset were matched with cell viability data from CTRP and Achilles datasets based on metadata (cell line, perturbation and concentration). It is possible to match one CTRP / Achilles cell viability instance with more than one LINCS-L1000 signature (same cell line and perturbation, but different perturbation time in LINCS-L1000). (B) Principal Component Analysis (PCA) of perturbation signatures from the CTRP-L1000 dataset. Each point represents a unique cell line - compound - concentration - perturbation time instance. Points are colored according to corresponding cell viability from CTRP screen (Spearman correlation between PC1 and cell viability: −0.260, p=1.28e-251). (C) Prediction of cell viability using linear models. Linear models were trained on CTRP-L1000 and Achilles-L1000 datasets (x axis). Prediction performance was evaluated on CTRP-L1000 and Achilles-L1000 datasets (color code) by calculating Pearson correlation (y axis) between predicted and observed values (results from 20 random sub-sampling validation, means +/- 95% CI). (D) The effects of perturbation time on the predictability of cell viability. Signatures from different time points (3, 6 and 24 hours for CTRP-L1000 and 96, 120 and 144 hours for Achilles-L1000) were used to train (x axis) and test (y axis) linear models. Baseline models were trained on cell line - perturbation ID data. Size and colors of circles are proportional with Pearson correlation, which is also labeled in selected cases (results from 20 random sub-sampling validations, means). (E) Gene Ontology enrichment of genes showing correlated expression with cell viability in the Achilles-L1000 dataset. GO terms associated with genes showing decreased (left) or increased (right) expression with increased cell death.

To explore the main factors behind the perturbation signatures, we performed Principal Component Analysis (PCA) on the signatures of the CTRP-L1000 dataset. While we observed no clustering of signatures in the first two principal component (PC) plane based on cell lines, perturbatogene compounds or perturbation time (SFig. 1), we found a clear relationship between PC1 and matched cell viability values (Fig. 1B, Spearman correlation: −0.260, p=1.28e-251).

Based on this PCA, we hypothesized that cell viability can be effectively predicted from perturbation signatures. We used linear models (*y=Xβ*) with L2 regularization trained on the CTRP-L1000 signatures (*X)* and cell viability values (*y*). Using random sub-sampling validation (Methods) we were able to predict “within” the CTRP-L1000 dataset (Fig. 1C) with average Pearson correlation 0.58 (predicted vs. observed cell viability, average log10(p)<-300) while the performance of “within” Achilles-L1000 prediction models was 0.49 (average log10(p)<-300). Furthermore, we were able to predict cell viability “across” the two dataset, predicting cell viability in the CTRP-L1000 dataset with models trained on Achilles-L1000 and vice versa. (Fig. 1C, average Pearson correlation: 0.32 and 0.17, average log10(p) values −206 and −273, respectively). This suggests that perturbation signatures are associated with cell death independent of the perturbation agent.

As previously mentioned, LINCS-L1000 dataset contains signatures from different elapsed time between perturbation and measurement. To analyse the effect of this elapsed time on the prediction performance, we split the CTRP-L1000 and Achilles-L1000 datasets based on measurement times (resulting CTRP-L1000-3h, CTRP-L1000-6h, CTRP-L1000-24h, Achilles-L1000-96h, Achilles-L1000-120h and Achilles-L1000-144h datasets, STable 1). We also introduced baseline models (linear models trained not on signature - cell viability data, but cell and perturbation ID - cell viability data, see Methods for further details) to benchmark our signature-based models prediction performance (Fig. 1D). We trained linear models (with L2 regularisation) for each time specific dataset, and tested them for the same dataset (“within” dataset prediction) and all the other datasets (“across” dataset prediction) using random sub-sampling validation. Baseline models were only used in the “within” dataset setting. While all signature-based models performed reasonably well in the “within” dataset setting (Fig. 1D diagonal), CTRP-L1000-24h and Achilles-L1000-96h models reached the best performance in the “across” dataset prediction. These two models also reached comparable performance (0.61 vs. 0.67 and 0.51 vs. 0.61 Pearson correlation, respectively) with the baseline models, and were able to make translatable predictions across CTRP and Achilles datasets (which would be impossible for the baseline models, as none of the perturbations are shared between these two datasets). Based on this benchmark, we selected CTRP-L1000-24h and Achilles-L1000-96h models for further use in this study. Effective performance of linear models across compound- and shRNA-based viability datasets suggests that there is some general transcriptomics signature of cell death. To further investigate this, we calculated Pearson correlation and significance between cell viability and gene expression on the Achilles-L1000-96h dataset for each gene, and performed Gene Ontology (GO) enrichment analysis based on these correlation and p values. The most significantly enriched GO terms were closely related to cell death and proliferation process (Fig. 1E). We obtained similar results using the CTRP-L1000-24h dataset (SFig 2), also suggesting the presence of perturbation independent cell death signature.

### 2.2 Signature of cell death as a confounding factor for mechanism of action discovery

One important application of perturbation signatures is mechanism of action (MoA) discovery (Iorio et al., 2010; Subramanian et al., 2017). Compounds having same / similar target molecules lead to similar transcriptomics response, which can help to identify previously unknown targets of compounds. However, as shown in the previous section, cell death can lead to a specific signature.

To analyze if this can be a confounding factor to infer mechanism of action, we took random pairs of samples from the CTRP-L1000-24h signatures, and calculated signature similarity using Spearman correlation between the signature vectors (see Methods for detailed description of sampling strategy). We focused on the the similarities between signatures of non-toxic (defined by cell viability>0.8 for the given cell line, drug and concentration) perturbations with shared MoA (based on (Corsello et al., 2017)), and highly toxic (cell viability<0.6) perturbations with different MoA (Fig. 2A). We sampled signatures irrespective of cell line or by sampling signature pairs from the same cell line to analyse the general and cell lines specific nature of signature similarity. Signature pairs of non-toxic perturbations with shared MoA are more similar than random pairs (Medians: 0.05 vs. 0.006 and 0.113 vs. 0.01, for cell line irrespective and same cell line sampling, respectively, Mann-Whitney U p values: 1.48e-108 and <1e-300). More interestingly, signature pairs of toxic perturbations with different MoA are also more similar than random pairs (Medians: 0.079 vs. 0.006 and 0.123 vs. 0.01, for cell line irrespective and same cell line sampling, respectively, Mann-Whitney U p values: 6.08e-281 and <1e-300) while similarity between toxic, different MoA signatures was higher / comparable with non-toxic, shared MoA signature similarity (Mann-Whitney U p values 1.54e-34 and 0.08 for cell line irrespective and same cell line sampling, respectively). These results suggest that cell death / toxicity is at least as important factor for signature similarity as mechanism of action, potentially confounding signature based MoA discovery.

**Figure 2 -.**
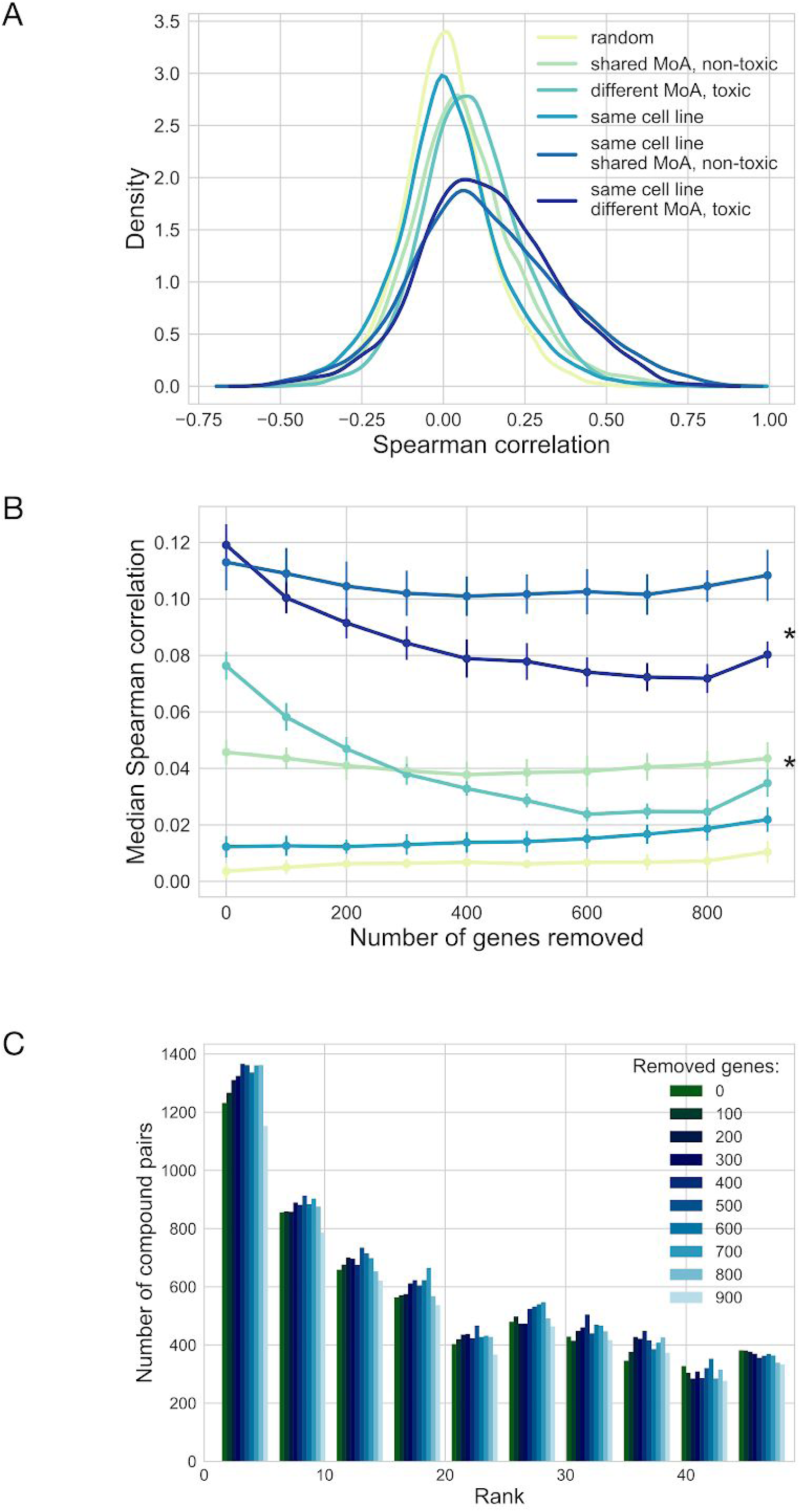
Cell death based signature as a confounding factor for mechanism of action discovery. (A) Mechanism of action and cell death based signature similarity. Random pairs of samples were taken from the CTRP-L1000-24h signatures with the following constraints (color code): absolutely random, non-toxic (cell viability>0.8) compounds with shared MoA and strongly toxic (cell viability<0.6) compounds with different MoA. Signature pairs were sampled independent of the cell line or from the same cell line. Spearman correlation (x axis) was used as signature similarity metric. Density plot shows results from 10,000 random samples / group. (B) Effect of removing cell death correlated genes on signature similarity. Top n (x axis) genes with highest absolute Pearson correlation with cell viability were removed from the signatures before similarity calculation (1,000 sample pairs repeated 10 times, mean +/- SD, *: p<2.2e-16 for number of removed genes, ANOVA). (C) Effect of removing cell death correlated genes on MoA discovery. Average signatures for 2485 compounds from LINCS-L1000-MoA dataset was calculated using MODZ method. For each compound, other compounds were ranked based on signature similarity (Spearman correlation). Histogram (ranks: 1-50) shows the ranks of compounds with shared MoA. Top n (color codes) genes with highest absolute Pearson correlation with cell viability were removed before consensus signature and signature similarity calculation (p=7.22e-73 from Kruskal-Wallis test).

To reduce this unwanted, cell death based signature similarity, we systematically removed the genes with highest absolute Pearson correlation with cell viability, and calculated signature similarity based on these reduced signature vectors (Fig. 2B). While removing these genes did not strongly affect the MoA based similarity (ANOVA p values: 0.23 and 0.09 for cell line irrespective and same cell line sampling, respectively, absolute effect sizes <1e-5/removed gene), it significantly reduced the similarity of toxic signatures (ANOVA p values <2.2e-16, absolute effect sizes >4e-5/removed gene). In contrast, removing genes randomly did not affect signature similarity (SFig. 3A, absolute effect sizes <1.5e-5/removed gene).

While this analysis suggests that removing genes correlated with cell viability can improve MoA discovery, the CTRP-L1000-24h dataset is biased toward toxic compounds (as in the CTRP screen anti-cancer drugs are used), so the effect of removing these genes is not so surprising. To analyse our method on a more balanced dataset, we selected all of the 24 hours compound perturbation signatures from LINCS-L1000 with known target molecule (LINCS-L1000-MoA dataset, 2485 compounds, STable 1) (Corsello et al., 2017). For each compound, we calculated consensus signatures (irrespective of cell line and concentration) using MODZ method (Moderated z-score, see in Methods), and calculated Spearman correlation as similarity metric between all pairs of average compound signatures. Following this, we calculated for each compound the ranks of other compounds with shared MoA based on average signature similarity, simulating the MoA identification process for an unknown compound (see Methods for more detailed description). Using reduced signatures significantly decreased these ranks (i.e. increased identification of compounds with shared MoA, Fig. 2C for ranks 1-50 and SFig. 3B for full rank distribution), having a top performance with 700 removed genes (median rank 964 vs. 1030 with full signature, Mann-Whitney U test p value: 1.36e-43). Removing random, not cell viability correlated genes did not affect the rank distribution (SFig. 3C) in this dataset either.

### 2.3 LINCS-L1000 as a cell viability assay

LINCS-L1000 contains a large number of chemical perturbations (that we shall call LINCS-L1000-Chem subset, 21921 compounds, STable 1), where most of the used compounds are not known anti-cancer drugs. We hypothesized that some of these drugs can have an unexpected, cell line specific anti-cancer activity that can be identified by predicting cell viability from the perturbation signatures. We used CTRP-L1000-24h and Achilles-L1000-96h models (based on their top performance, Fig. 1D) to predict cell viability for the whole LINCS-L1000-Chem dataset and identified several known and also potentially clinically interesting drugs with cell line specific toxicity.

To further evaluate the prediction performance of these linear models, we compared the predicted cell viability values with the results of NCI60 screen (Shoemaker, 2006). As NCI60 is a discovery screen (most of the drugs used did not have anti-cancer activity, SFig. 4A-C) it is a realistic benchmark dataset for our LINCS-L1000-Chem predictions. We found an intersection of 583 compounds and 6 cell lines (NCI60-L1000-24h dataset, STable 1) between the two screens. We binarized GI50 (50% growth inhibition) results of the NCI60 screen (effective / ineffective anti-cancer drugs, where ineffective means 50% growth inhibition was not reached in the used concentration range, see Methods for further details), and compared them with the predicted cell viability (lowest value for each compound - cell line pair) with ROC curves. We also selected the intersection between LINCS-L1000, NCI60 and CTRP datasets (NCI60-CTRP-L1000-24h dataset, 99 compounds, STable 1), where we could compare the performance of the linear models against a “ground truth based” method. In this case CTRP drug sensitivity metric (area under the dose response curve, AUC) was used to predict the effectiveness in NCI60. ROC analysis revealed that the performance of the perturbation signature based models is comparable with the “ground truth based” method (adjusted p values for AUC difference > 0.05), and that Achilles-L1000-96h model reached better performance than CTRP-L1000-24h model on the whole NCI-L1000-24h dataset (AUC=0.78 vs. 0.71, p=1.398e-15). We had similar results when used TGI (total growth inhibition) or LC50 (50% lethal concentration) as NCI60 drug sensitivity metric (SFig. 4D and E), further suggesting the reliable performance of our models to predict cell viability even in independent datasets.

As Achilles-L1000-96h model had the best performance across all benchmarking experiments (Fig. 1D, Fig. 3A, SFig. 4D and E) we analysed the predictions of this model for the whole LINCS-L1000-Chem dataset (Supplementary File 1). We focused on the highest concentration instances of each compound, and selected the lowest (most toxic) and highest (less toxic / most proliferative) predicted cell viability for each compound. These lowest and highest predicted cell viability instances were coming from different cell lines, thus allowed us to analyse the general and cell-specific compound toxicity together. We plotted lowest and highest predicted cell viability values for each compound against each other, and grouped compounds to toxic and proliferative groups based on arbitrary thresholds (5th and 95th percentile of cell viability values from Achilles-L1000 data) of predicted cell viabilities (Fig. 3B). Drugs from the CTRP screen (“known anti-cancer drugs”) had a higher representation in the toxic group (50.8 percent vs. 7.1 percent for all compounds, Fisher exact test p value: 1.49e-98). In the general toxic group (toxic effect in all screened cell lines) we found proteasome inhibitors (delanzomib, oprozomib), detergent (benzethonium), cell cycle inhibitor (PHA-848125), topoisomerase inhibitors (teniposide, SN-38), anti-eukaryotic antibiotics (pyrvinium, ivermectin, niclosamide) and plant derivatives (bruceantin, cucurbitacin-i, homoharringtonine). Our predictions also revealed the known proliferative effect of epithelial growth factor receptor (EGFR) agonist ligands (EGF, TGFa and betacellulin) in breast cancer cell lines. More interestingly, we identified several cyclin-dependent kinase (CDK) inhibitors (dinaciclib, CGP-60474, PHA-767491) with cell line specific predicted proliferative / toxic effect.

**Figure 3 -.**
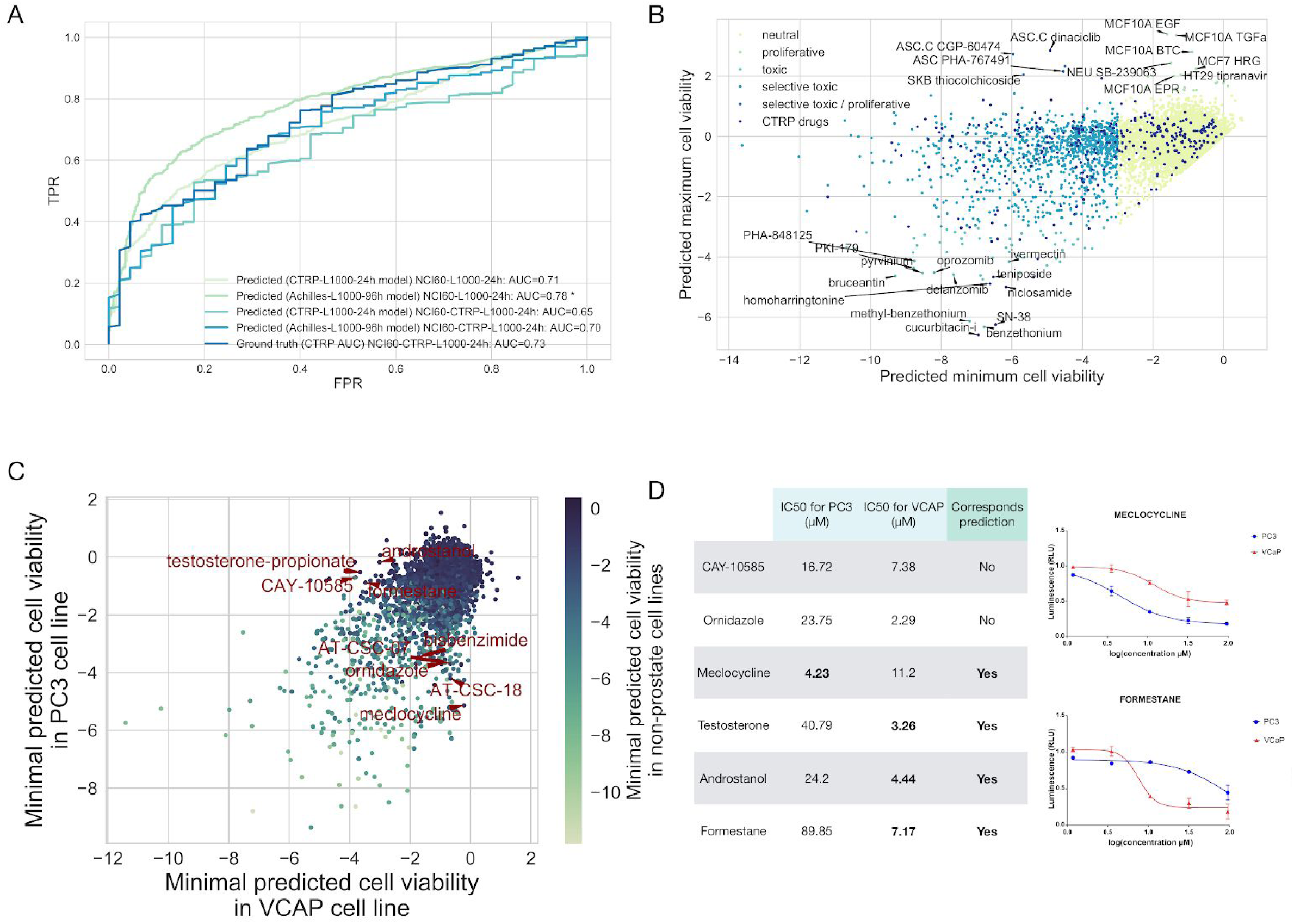
Predictions of cell viability for the whole LINCS-L1000 dataset. (A) ROC analysis of the prediction performance of linear models on NCI60 data. Cell viability was predicted for the intersection of NCI60 and LINCS-L1000 or for the intersection of NCI60, CTRP and LINCS-L1000 datasets (NCI60-L1000-24h and NCI60-CTRP-L1000-24h respectively) using linear models trained on CTRP-L1000-24h or Achilles-L1000-96h data. Either these predicted cell viability values or the known AUC values from CTRP screen (CTRP AUC) were used to predict the binarised (effective / ineffective in the investigated concentration range) GI50 from NCI60 (*: p<1e-5 for difference between AUCs for CTRP-L1000-24h and Achilles-L1000-96h). (B) Classification of the compounds from the LINCS-L1000 dataset based on their effect on cell viability. Cell viability was predicted for the LINCS-L1000-Chem dataset (24 h signatures, highest used concentration) using Achilles-L1000-96h model. The minimum (x axis) and maximum (y axis) of predicted cell viability was plotted for each compound. Compounds were classified as toxic (predicted cell viability < -3) or proliferative (predicted cell viability >1.5) and colored based on toxicity and selectivity (based on maximal and minimal predicted value). Compounds present in CTRP dataset (known anti-cancer drugs) were also labelled. For selected compounds the name of the compound and the corresponding cell line is text labelled. (C) Cell selective toxicity of LINCS-L1000 compounds in prostate cancer cell lines. Minimal predicted cell viability for VCaP (x axis) and PC3 (y axis) prostate cancer cell lines was plotted for each compounds. Data points are color coded based on the minimal predicted cell viability in non-prostate cancer cell lines. For selected compounds showing selective toxicity in prostate cancer cell lines the name of the compound is text labelled. (D) IC50 values of experimentally validated compounds for VCaP and PC3 cell lines. For meclocycline and formestane dose response curves are also shown (right up and down, respectively), while other full dose response curves for other compounds are shown in SFig. 5.

To further analyse the ability of Achille-L1000-96h model to predict cell line specific compound toxicity, we focused on the two prostate cancer cell lines (PC3 and VCaP) present in the core cell lines of LINCS-L1000 (Fig. 3C). Our model predicted several compounds with cell line specific toxicity for these two prostate cancer cell lines, including HIF1A inhibitor CAY-10585 and several androgen receptor related compounds (androstanol, testosterone-propionate and formestane) for VCaP and antibiotics ornidazole and meclocycline for PC3 (Fig. 3C, text labelled data points). For these six drugs, with markedly different predicted toxicity in the two prostate cancer cell lines, we performed cell viability measurements (Methods). Half inhibitory concentrations (IC50s, Fig. 3D and SFig. 5) showed increased sensitivity of PC3 for meclocycline (4.23 μM vs. 11.2 μM for VCaP) and increased sensitivity of VCaP for testosterone, androstanol and formestane (3.26, 4.44 and 7.17 μM vs. 40.79, 24.2 and 89.85 for PC3, respectively), confirming 4 of our 6 predictions. In case of the two other predictions we observed low toxicity (ornidazole) and ambiguous results (lower IC50 but also lower maximal toxicity in VCaP cell lines for CAY-10585).

### 2.4 Drug perturbation signatures as features for machine learning models

As shown in the previous sections, cell viability can be effectively predicted from perturbation transcriptomics data. However, for these predictions, linear models already needed measurement of the perturbation signature of the investigated cell line - compound pairs. We then asked if it would be also possible to predict cell viability / drug sensitivity for cancer cell lines where the actual perturbation measurement was not performed - a much more challenging task. We reasoned that this could be attempted using drug specific consensus signatures together with cell line specific information to predict drug sensitivity.

To achieve this prediction of cell line specific drug sensitivity from consensus signature we used machine learning models and applied them to an independent data-set. We chose the GDCS (Iorio et al., 2016) data as the largest pharmacogenomic drug screening available. Before evaluating the performance of machine learning models, we analysed how the consensus drug signatures correspond to the known mechanism of action of GDSC drugs. Consensus signatures (MODZ method) were generated for GDSC drugs present in the LINCS-L1000 screen (GDSC-L1000-24h dataset, 148 drugs, STable 1). Relationship between consensus signatures were visualised after dimensionality reduction by t-SNE algorithm (Maaten and Hinton, 2008) (Fig. 4A). Some drugs with shared mechanism of action (e.g. MAPK, PI3K and HDAC inhibitors) formed clusters in t-SNE plane and, even more interestingly, we could observe several clusters formed by unrelated drugs. For example GSK3 inhibitors CHIR-99021 and SB216763 formed a cluster with PKC inhibitor Enzastaurin, MAPK7 inhibitors XMD8-92 and XMD8-85 formed a cluster with PLK inhibitor BI-2536 and JAK2 inhibitor Fedratinib while topoisomerase inhibitor Doxorubicin formed clusters with CDK inhibitors AT-7519 and CGP-60474 (Fig. 4A, inserts), which suggests consensus signatures can potentially reveal unknown similarities between anti-cancer drugs.

**Figure 4 -.**
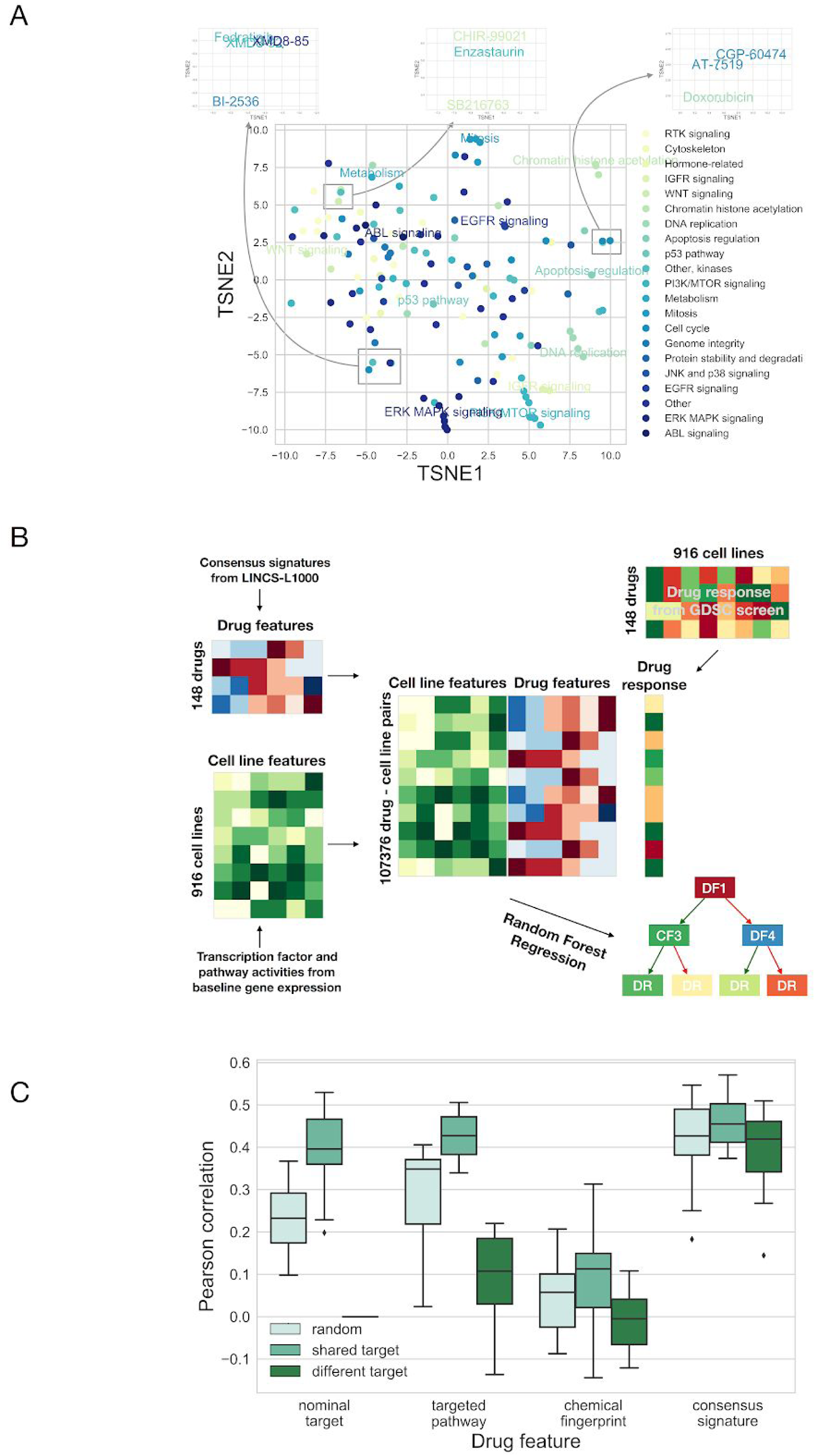
Consensus perturbation signatures for drug sensitivity prediction. (A) t-SNE plot (learning rate 80, perplexity 20, number of iterations 1000) of the consensus signatures of GDSC drugs. Data points (GDSC drugs) are colored based on targeted pathway. For selected clusters the targeted pathway is also text labeled. Inserts: selected clusters of drugs with different pathway annotation. (B) Schematic representation of machine learning pipeline. Cell line specific (histology, pathway and transcription factor activity) and drug specific (nominal target, targeted pathway, chemical structure based Extended-Connectivity Fingerprints and consensus signature) feature matrices were created for the cell lines and drugs of the GDSC study. Features were concatenated for each drug - cell line pair, where drug response was available in the GDSC screen and this feature matrix was used to train Random Forest Regression models. (C) Results of the machine learning models. Data was splitted into training and test set based on drugs (50-50% percent). Splitting was performed 3 different way (color code): randomly, or with constraint that for each drug in test set there was a drug with same nominal target in the training set (shared target), or with constraint that for each drug in test set there was no other drug with shared nominal target in training set (different target). Different drug specific features (x axis) was used by the models. Cell wise average Pearson correlation values are shown as boxplots for the different drug specific features / splitting strategies (results from 20 random sub-sampling validation).

We than used Random Forest Regression in multi-task setting (predicting drug sensitivity for different cell lines and drugs with the same model) to predict drug sensitivity (area under the dose response curve, AUC) values from the GDSC, using consensus perturbation transcriptomics signatures from the LINCS-L1000 study as features. To predict drug sensitivity in different cell lines and for different drugs, the Random Forest Regression model required cell line and drug specific features (Fig. 4B). As cell line specific features we used histology type, pathway (Schubert et al., 2018) and transcription factor (Garcia-Alonso et al., 2018) activities inferred from baseline gene expression (see Methods for further details), while consensus perturbation signatures were used as drug specific features. For benchmarking we also used other types of drug specific features (see Methods for further details): nominal target of the drug, targeted pathway of the drug and chemical structure based Extended-Connectivity Fingerprints (ECFP-like), to compare the performance of signature based features with them.

We focused on the prediction of drug sensitivity for new (for the model unknown) drugs. To do this, we splitted the GDSC dataset in two halves, based on the used drug (i.e. half of the drugs were in our training, other half in the test set). We used three different splitting schemes: random, shared target and different target. In shared target setting each drug in the test had a pair in the training set with the same nominal target, while in different target setting the drugs in training and test set had strictly different nominal targets (see Methods for further details). We evaluated prediction performance by calculating Pearson correlation between predicted and observed drug sensitivity (dose response AUC) values for each cell line (Fig. 4C). In the shared target setting nominal target, targeted pathway and consensus signature based features had similar performance (mean Pearson correlations: 0.40, 0.42 and 0.46 respectively, ANOVA p value for used model: 0.028). Not surprisingly nominal target based drug features were not useful in the case of different target sampling (mean Pearson r: 1.52e-18, p value from one sample t-test with 0 population mean: 0.79), while consensus signature based features outperformed targeted pathway features in this case (mean Pearson correlations: 0.39 and 0.09 respectively, p value from paired t-test: 1.1e-7). In summary, consensus signature based drug features for machine learning models had better performance than current gold standard drug specific features like nominal target or target pathway, and allowed cell line specific prediction of drug sensitivity.

## 3. Discussion

In this paper we integrated recent large-scale functional and pharmacogenomic studies to analyse the effect of cell viability on perturbation transcriptomics signatures. We found that cell viability - cell death and cell proliferation - has a major contribution to the perturbation signatures. While this association between cell viability phenotype and transcriptional signatures enables efficient prediction of cell viability values from perturbation signatures, it can also mask the compound specific transcriptional signal, thus confounding discovery of mechanism of action.

Using perturbation metadata from the LINCS-L1000, CTRP and Achilles projects, we composed the (to our knowledge) largest matched cell viability - perturbation signature dataset with more than 15,000 compound and more than 75,000 shRNA treatments. Principal Component Analysis the CTRP-L1000 dataset revealed that the first PC (explaining 9% of total gene expression variance) is associated (Fig. 1B) with cell viability. The cell line and perturbation independent nature of PC1 was already described in one of the original LINCS-L1000 manuscripts (Smith et al., 2017), and it was speculated that it is connected to some general biologic effect, but it was not explicitly analyzed previously. Also, a recent analysis (Jung et al., 2018) compared perturbation signatures with cell viability. Jung et al. matched perturbation signatures from the LINCS-L1000 screen with corresponding cell growth inhibition (cell viability) values from an earlier version of the GDCS with 639 cell line - compound pairs (Garnett et al., 2012) and used these data identify essential gene signatures, while our study focused on the predictive value of perturbation signatures and the confounding effect of cell death on signature similarity and mechanism of action identification.

Based on this association between cell viability and perturbation signatures, we were able to predict cell viability from the gene expression data. Most importantly, linear models trained on drug perturbations (CTRP data) were able to predict cell viability after shRNA treatment (Achilles data), and vice versa (Fig. 1C and 1D). While baseline models - using perturbation and cell line information - were also able to reach effective performance within a given dataset (CTRP or Achilles), this effective across dataset prediction is unique for the perturbation signature based models. Also several studies investigated the predictability of drug sensitivity (Ali and Aittokallio, 2018; Costello et al., 2014; Iorio et al., 2016) and gene essentiality (Gönen et al., 2017) with good performance, but translatable prediction was not attempted between these different, but related phenotypes. There could be two main reasons for the effective across dataset prediction performance of our methods. First, that models can learn the drug / shRNA specific changes in signatures, and utilize the similarities between signatures to predict cell viability across different perturbations. The second possibility is that there is a specific cell death signature, independent of the original perturbation agent, and linear models can learn this signature. Our Gene Ontology analysis (Fig. 1E) suggests the latter one. We also analyzed how the elapsed time between perturbation and transcriptomics measurement affects the predictability of cell viability. While the two best performing models (CTRP-L1000-24h and Achilles-L1000-96h) have the largest amount of data available (STable 1), the poor performance of 3 and 6 hour models also suggest that the cell viability related gene expression changes are only observable after longer perturbations. Importantly, our models were trained on transcriptomics data from the LINCS project and cell viability data form the CTRP and Achilles project and were also evaluated on the NCI60 cell viability data (Fig. 3A). Hence, we used data from 4 different sources. The effective performance of the models across these different studies suggests the underlying biological phenomenon is robust, and also provides a step to help address the challenge of translating models across drug sensitivity screens (Cancer Cell Line Encyclopedia Consortium and Genomics of Drug Sensitivity in Cancer Consortium, 2015; Haibe-Kains et al., 2013).

Using these models we were able to predict cell viability for the whole compound perturbation part of the LINCS-L1000 study. We were able to identify not only well known general toxic compounds like detergents, proteasome or topoisomerase inhibitors, but also compounds leading to proliferation like EGF in breast cancer cell lines (Fig. 3B). Most interestingly, several CDK inhibitors had cell line specific toxic or proliferative effect, where proliferative effect was observed in adipocyte stem cells (ASCs). While CDK inhibitors can uncouple cell cycle form apoptotic function (Le et al., 2005), so this observation can be a false positive, some experimental evidence also suggests that CDKs can have paradox proliferative effect in stem cells (Li et al., 2012). We also analysed cell line specific predictions in prostate cancer cell lines VCaP and PC3, and performed experimental validations for 6 compounds showing marked difference of toxicity between these two cell lines. We found that several androgen receptor signaling related compounds (like androstanol and testosterone-propionate) have selective toxic effect in VCaP cell lines. This paradoxical toxic effect of androgen receptor agonists have been also reported in another castration resistant prostate cancer model (Nakata et al., 2016). Our results show that sensitivity of different prostate cancer cell lines can be markedly different for androgen treatment, also suggesting androgens as therapeutic options in selected cases of metastatic disease (Thelen et al., 2013). We also observed selective toxicity of meclocycline in PC3 cell lines. As this drug is an antibacterial antibiotic, it could be also used as potential treatment with low adverse effect profile in prostate cancer. Interestingly, meclocycline is a tetracycline antibiotic, a group of drugs known to have a metalloprotease inhibitor effect (Saikali and Singh, 2003). Matrix metalloproteinases are recently proposed as important molecules and drug targets in prostate cancer (Gong et al., 2014).

While our models were able to predict cell viability from the actual measured perturbation signatures, one can argue that in this case the experiment is already performed, and if somebody is interested for cell viability, testing viability is simpler and cheaper than perturbation transcriptomics profiling. Yet, our machine learning predictions on the GDSC dataset show that using consensus signatures (delivered from a small number of core cell lines) as features for machine learning models allows the prediction of drug sensitivity also in new cell lines. Our results show that consensus signature outperforms gold standard features like nominal target, targeted pathway and chemical fingerprints (Fig. 4C) and allows prediction of drug sensitivity for new drugs (unrelated to the ones present in the training set). Also our models can be used to predict cell viability in previously performed gene expression studies, where cell viability data was not measured.

Clustering of the GDSC drugs based on the consensus signatures (Fig. 4A) also revealed important information regarding their mechanism of action. While some drugs with the same MoA (e.g. ERK/MAPK inhibitors and PI3K/MTOR inhibitors) formed tight clusters, we also observed clusters of apparently unrelated drugs. The similarity of PKC inhibitor Enzastaurin signature to GSK inhibitors have been reported in the original LINCS-L1000 study (Subramanian et al., 2017), while the connection between CDK inhibitors and Doxorubicin have been also described previously (Iorio et al., 2010). We also observed a cluster composed of XMD8-92, XMD8-85, BI-2536 and Fedratinib (MAPK7, MAPK7, PLK and JAK2 inhibitors, respectively). Recent experiments support that all of these drugs have a common BET inhibitor effect (Ciceri et al., 2014; Lin et al., 2016), probably responsible for the signature similarity.

While the association of perturbation signatures with cell viability enables effective prediction, it can also be a confounding factor for mechanism of action discovery. The similarity between toxic compounds with different MoA is comparable with the similarity between non-toxic compounds with the same MoA (Fig. 2A) which process can also negatively influence MoA discovery (Fig. 2C). Importantly, removing genes with high absolute correlation with cell viability helps to overcome this problem (Fig. 2B and 2C), and can help to analyse the results of perturbation transcriptomics signatures more rigorously.

The linear models and pre-calculated predicted cell viability values (Supplementary File 1) can be an important resource for further studies working with perturbation gene expression signatures. Also, our results suggest that removing cell death correlating genes from the gene expression signature can help to better interpret the similarities between signatures and identify mechanism of action. As drug sensitivity prediction with machine learning models is an important area of current research, we think our work gives an important new feature, consensus perturbation signatures, to this field. Finally, while we analysed solely cell viability as cellular phenotype, the methods presented here possibly can be used also in the context of other perturbation - phenotypic measurement studies.

## 4. Methods

### 4.1 Databases and data preprocessing

We used the Phase I and Phase II LINCS-L1000 perturbational profiles (Subramanian et al., 2017) (GSE92472 and GSE70138 respectively, downloaded from Gene Expression Omnibus (Barrett et al., 2013)) in this study. Replicate-collapsed differential expression signatures (Level5 dataset) of the measured (landmark) genes were used in our analysis pipeline. For accessing L1000 signatures, we used cmapPy Python library (Enache et al., 2018). Phase I and Phase II data were merged, and signatures corresponding to the same conditions (treatment, cell line, time and concentration in case of compounds, or treatment, cell line and time in case of shRNA) were averaged using the MODZ method.

CTRPv2 cell viability dataset (Seashore-Ludlow et al., 2015) was downloaded from CTD^2^ Data Portal (https://ocg.cancer.gov/programs/ctd2/data-portal). For further analysis the post-quality-control cell viability values were used. We matched CTRP and L1000 instances based on cell line and Broad compound IDs. For matching compound concentrations we matched the instances between CTRP and L1000, where the concentration difference was the smallest, and the absolute log10 concentration difference was smaller than 0.2 (~1.5 fold concentration difference).

Achilles 2.4.6 and 2.19.2 datasets (Tsherniak et al., 2017) were downloaded from Project Achilles Data Portal (https://portals.broadinstitute.org/achilles). We used the shRNA log fold change scores in our analysis (i.e. without separating on- and off-target effects of shRNAs). We matched Achilles and L1000 instances based on cell lines and shRNA treatment. As LINCS-L1000 identifies shRNAs with Construct ID and Achilles uses Barcode Sequence, we mapped these two identifiers with the help of the reference files from the Genetic Perturbation Platform (https://portals.broadinstitute.org/gpp/public/).

NCI60 drug toxicity datasets (Shoemaker, 2006) (GI50, LC50 and TGI values) were downloaded from the Developmental Therapeutics Program data portal (https://dtp.cancer.gov/discovery_development/nci-60/). We restricted our analysis to those compounds that overlap between L1000 and NCI60 screens. For easier comparison, we extracted the PubChem Compound IDs using PUG REST services in R (https://pubchemdocs.ncbi.nlm.nih.gov/pug-rest-tutorial). For compounds in the NCI60 dataset, we converted Substance IDs to Compound IDs. Whereas, for compounds in the L1000 dataset, we either used the Compound IDs directly (when available) or retrieved them based on the InChi keys. As the original (PubChem Compound ID based) intersection between L1000 and NCI60 datasets were relatively small (373 compounds), we retrieved the name synonyms of each compound (using PubChemPy Python library) and matched compounds based on their name synonyms that resulted in 583 shared compounds between the two datasets.

### 4.2 Moderated z-score (MODZ)

We calculated consensus signatures using a moderated z-score, described in the original LINCS-L1000 paper (Smith et al., 2017; Subramanian et al., 2017). Basically, for a set of signatures pairwise Spearman correlation matrix was calculated. Diagonals (self correlations) were set to 0, while negative correlations were set to a small value (0.01). Weight for each signature was the sum of these correlation values row-wise (normalised so, that sum of weights was 1). The final consensus signature was calculated as a weighted average of the signatures.

### 4.3 Linear models

We used linear regression (y=Xβ) with L2 regularization (alpha=1.0) to predict cell viability (y, n*1 column vector, where n is the number of samples) from perturbation gene expression signatures (X, n*g matrix, where n is the number of samples, g is the number of genes in signatures). To evaluate prediction performance, we used random sub-sampling validation strategy: half of a given dataset was used to train the models, while cell viability was predicted for the other half of the dataset. This process was repeated 20 times and we used Pearson correlation between the predicted and observed cell viability values, as an evaluation metric. We call “within dataset prediction”, when the training and the test data come from the same dataset (e.g.: CTRP-L1000-24h). In contrast, when we train a model on one dataset, and predict cell viability for an other dataset (e.g.: CTRP-L1000-24h and Achilles-L1000-96h, respectively), we use the term “across dataset prediction”. In case of across dataset prediction, we trained the linear model on half of the training data, but evaluated it on the whole test data. For baseline model, feature matrix (X) was composed of indicator vectors for cell lines and perturbations and a vector containing log10 drug concentration (only in case of CTRP-L1000 datasets).

### 4.4 Gene Ontology Enrichment

We calculated Pearson correlation r and p values between cell viability and gene expression for each gene in the Achilles-L1000 and CTRP-L1000 datasets. We used these r and p values as input for piano R package (Väremo et al., 2013), and calculated Gene Ontology enrichment (mean Gene Set Analysis method). We report FDR adjusted p values for the top 10 Gene Ontology term.

### 4.5 Average silhouette analysis

For evaluation of the different factor (cell line, drug, perturbation time, cell viability) based clustering of CTRP-L1000 data points in the first two principal component plane, we used average silhouette analysis with Euclidean distance. Silhouette coefficient (b-a) / max(a,b) was calculated for each datapoint, where a is mean intra- and b is mean nearest-cluster distance. For each clustering factor the average of Silhouette Coefficients were calculated (scikit-learn Python library (Pedregosa et al., 2011)). In this case negative average silhouette score corresponds to the absence of clustering, while positive score means clustering of data points based on the selected factor.

### 4.6 Signature similarity analysis

We analysed mechanism of action and cell death based signature similarity using Spearman correlation as similarity metric. We took random samples from the CTRP-L1000-24h dataset, where signature pairs were coming from non-toxic (cell viability>0.8) perturbations with shared MoA, or strongly toxic (cell viability<0.6) perturbations with different MoA. For MoA definition we used compound metadata (gene symbol of protein target, target or activity of compound) from the CTRP screen. For defining non-toxic and strongly toxic cell viability thresholds, we fitted Gaussian Mixed model on cell viability values (mean of non-toxic group ~1.0, SD ~0.1, so thresholds 0.8 and 0.6 corresponds to mean - 2*SD and 4*SD respectively). We performed cell line irrespective sampling and also sampled signature pairs from the same cell line. For analysing the effect of reduced signatures, we either removed random n genes from the perturbation signatures or the top n genes with highest absolute Pearson correlation with cell viability phenotype. To prevent “data leakage”, Pearson correlation between gene expression and cell viability was calculated from the Achilles-L1000-96h dataset.

For analysing the signature similarity for a larger part of the LINCS-L1000 dataset, we selected compounds with known MoA based (Corsello et al., 2017) on the Drug Repurposing Hub (clue.io/repurposing, May 2018 version). We calculated consensus signature for each compound using the MODZ method, using all signatures for the given compound. For each compound, we ranked other compounds based on signature similarity (Spearman correlation between signature vectors) and collected ranks for compounds with the same MoA. For reduced signatures, genes were removed from the signatures before consensus signature calculation.

### 4.7 NCI60 validation

NCI60 screen (Shoemaker, 2006) calculated GI50 (50% growth inhibition concentration), TGI (total growth inhibition concentration) and LC50 (50% lethal concentration) as drug sensitivity metrics for the used cell line - compound pairs. A sensitivity metric was only given, if the effect (50% growth inhibition etc.) is reached in the used concentration range, otherwise the maximal tested drug concentration is indicated. Based on this we defined the delta concentration metric: sensitivity metric - maximal tested concentration (log10 concentration values). Based on this definition of delta concentration values, delta concentration < 0 means effective drug, so we used this threshold for binarization for ROC analysis. For ROC analysis we used binarized delta concentration values as true positive / negative, while the predictions of linear models were used as target scores. For each cell line - compound pair we used the lowest predicted cell viability (when multiple signatures were available) as target score. For the “ground truth” model (predicting drug sensitivity in NCI60 from drug sensitivity in CTRP for shared cell lines and compounds) the dose response AUC values from CTRP screen were used.

### 4.8 Cell viability assay

PC3 and VCaP cell lines were purchased from ATCC. Cytotoxicity of different test compounds was studied on both PC3 and VCaP cell lines by determining the number of viable cells based on quantitation of ATP using the CellTiter-Glo® Luminescent Cell Viability Assay (Promega, Mannheim, Germany).

Cells were seeded into 384-well plate (Corning Life Sciences, Tewksbury, MA, USA) at 1.000 cells/well density in 40 μl media and incubated for 4h at 37 °C. Test compounds were dissolved in dimethyl sulfoxide (DMSO, Sigma, Budapest, Hungary) and cells were treated with an increasing concentration of drugs (1, 11 μM to 90 μM). The highest applied DMSO content of the treated cells was 0.5%. After 48h incubation at 37°C under 5% CO2, 40 μl CellTiter-Glo® Reagent (Promega) were added to each well and the luminescent signal was recorded by luminometer (VICTOR Multilabel Plate Reader, Perkin Elmer). Viability was calculated with relation to untreated control cells after extracting signals from blank wells containing only culture medium. IC50 values (50% inhibiting concentration) were calculated by GraphPad Prism®5 (La Jolla, CA, USA).

### 4.9 Machine learning models

Transcription factor and pathway activities as cell line specific features were calculated from baseline gene expression data (Iorio et al., 2016). For each g gene and c cell line standardised gene expression was calculated as Z_gc_=(E_gc_-μ_g_)/σ_g_ where E_gc_ is gene expression, μ_g_ and σ_g_ are means and standard deviations of a gene across cell lines. From these standardised gene expression values transcription and pathway activities were calculated using DoRothEA (Garcia-Alonso et al., 2018) and PROGENy (Schubert et al., 2018) methods, as described previously.

Nominal target and targeted pathway features were created from manually annotated drug metadata from the GDSC portal (www.cancerrxgene.org). Extended-Connectivity Fingerprints (ECFP-like) were generated by using the RDKit fingerprint module in KNIME with the radius and number of bits being set to 2 and 256, respectively. For consensus signature based features we mapped the PubChem compound IDs of the GDSC drugs with that of the compound IDs in LINCS-L1000 dataset. For each GDSC drug a consensus signature was calculated by using the MODZ method (using all available 24 hour signatures, irrespective of cell line and concentration). To reduce the dimensionality (978) of these signature features, we performed PCA and selected the first 40 Principal Components (explained variance: 95%).

We used Random Forest Regression models (with 50 trees) from scikit-learn Python library (Pedregosa et al., 2011). For each model the specified drug features (targets, pathways, chemical fingerprints or consensus signatures) and all cell-specific features (histology, pathway and transcription factor activities) were concatenated to create the feature matrix. Area under the dose response curve (AUC) was used as drug sensitivity metric. We used a random sub-sampling strategy to train and evaluate model performance. For each run, half of the drugs were included in the training set, while the remaining half formed the test set. 3 different methods were used to split the GDSC drugs into training and test set: random splitting, splitting where for each drug in the test set there was a corresponding drug with shared target in the training set and splitting where all drugs targeting a given protein were either in the test or the training set. For evaluation, Pearson correlation was calculated for each cell line between the predicted and observed AUC values, and these cell wise Pearson correlations were averaged. This random sub-sampling validation process was repeated 20 times.

### 4.10 Statistical analysis

Statistical significances were calculated using the corresponding functions of SciPy library (Pearson correlation, Spearman correlation, Mann-Whitney U test, Kruskal-Wallis H test, Fisher exact test, paired t-test) and ANOVA and pROC (Robin et al., 2011) from R.

## Supporting information

## 5. Acknowledgements

This work was supported by European Union Horizon 2020 research and innovation programme under grant agreement No 668858. B.Sz. was supported by the Premium Postdoctoral Fellowship Program of the Hungarian Academy of Sciences. We would like to thank Luis Tobalina Segura, Javier Perales Patón, Mi Yang, Michael Schubert and Francesco Iorio for reading the manuscript and providing useful comments.

## 6. Authors contributions

BS designed the research, performed all analysis and wrote the manuscript. VS performed ligand matching from databases and contributed to results interpretation and manuscript writing. RA performed and analysed validation experiments, supervised by LGP. JSR supervised the project and contributed to results interpretation and manuscript writing.

## 7. Conflicts of interest

The authors declare no conflict of interest.

## Supplementary Materials

### Supplementary Files

#### Supplementary File 1

Supplementary File 1 contains LINCS-L1000 signature IDs with matched cell viabilities from Achilles and CTRP screens, coefficients of the CTRP-L1000–24h and Achiles-L1000–96h models and LINCS-L1000 signature IDs with predicted cell viabilities for the LINCS-L1000-Chem dataset.

**Supplementary Figure 1 -.**
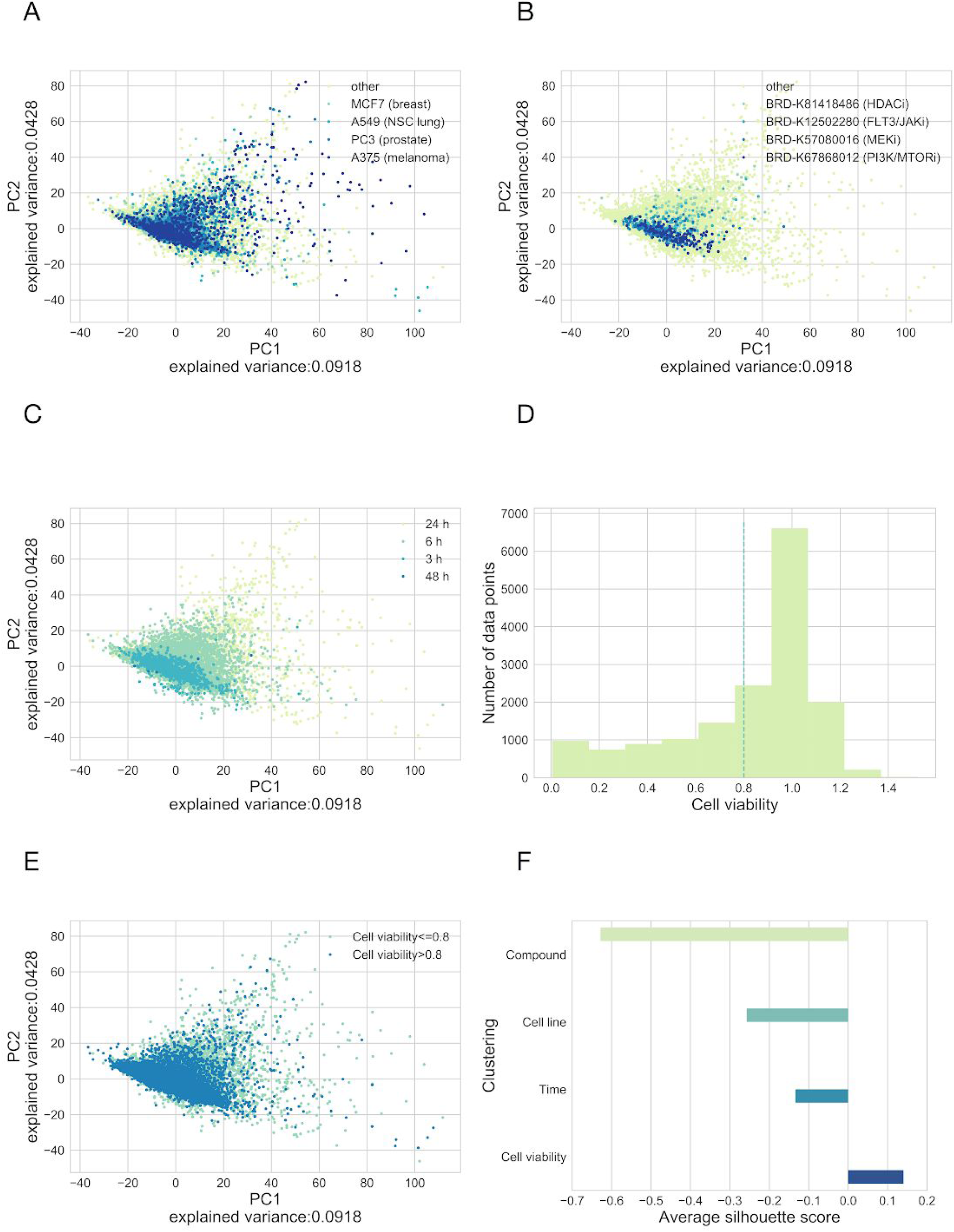
Clustering of L1000 signatures based on different factors. Principal Component Analysis (PCA) was performed on the perturbation signatures from CTRP-L1000 dataset. Each point represents an unique cell line - compound - concentration - perturbation time instance. Points are colored according to cell lines (A), compound used for perturbation (B), perturbation time (C) and cell viability (E). Only selected compounds and cell lines (with largest number of data points) are color labelled. For cell viability based clustering we selected 0.8 as threshold for toxic / non-toxic cluster based on the histogram (D) of cell viability values (~2 SD below mean based on Gaussian Mixed model). *We* performed average silhouette analysis using the different clustering factors (F).

**Supplementary Figure 2 -.**
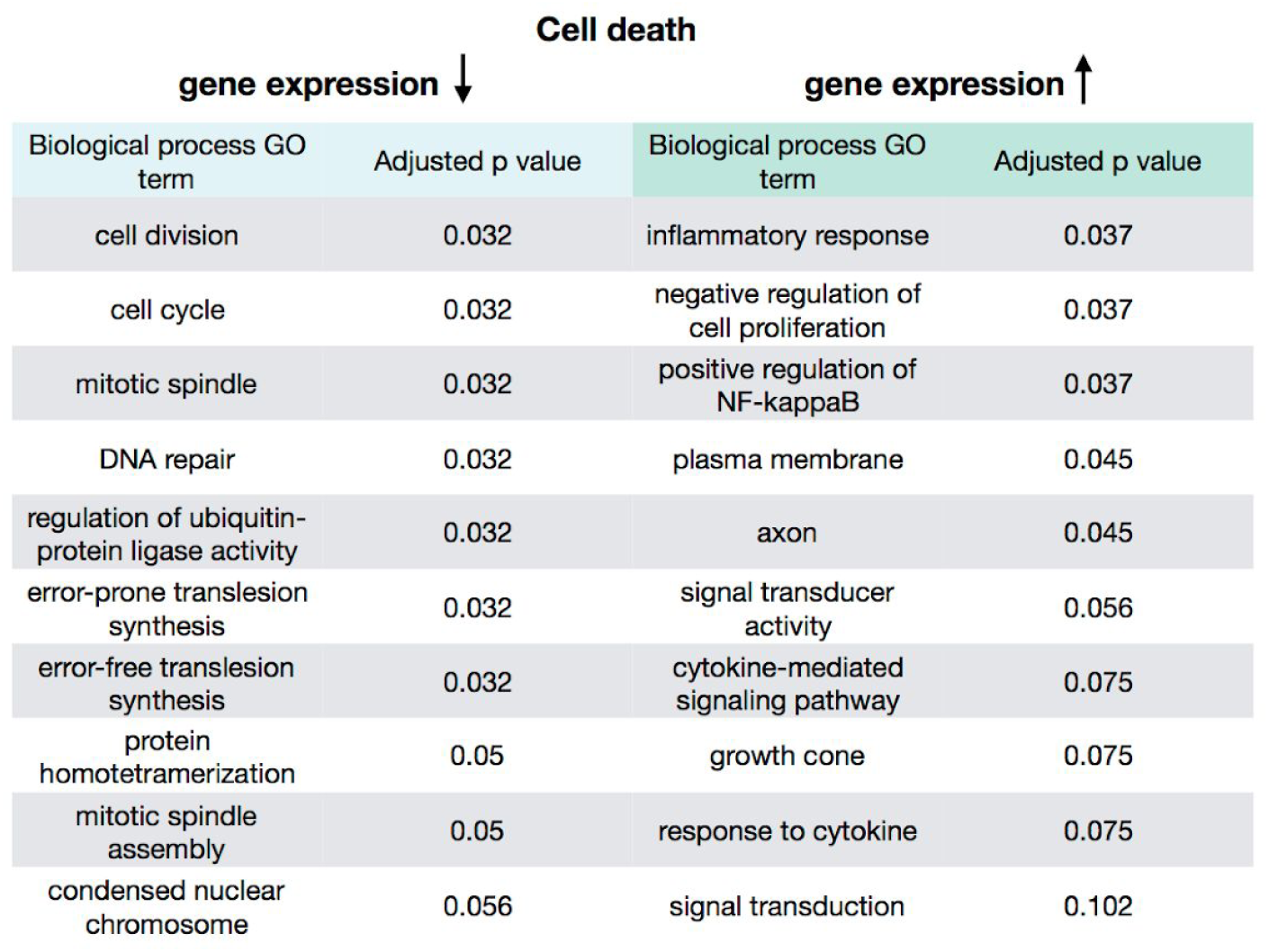
Gene Ontology enrichment of genes showing correlated expression with cell viability in the CTRP-L1000 dataset

**Supplementary Figure 3 -.**
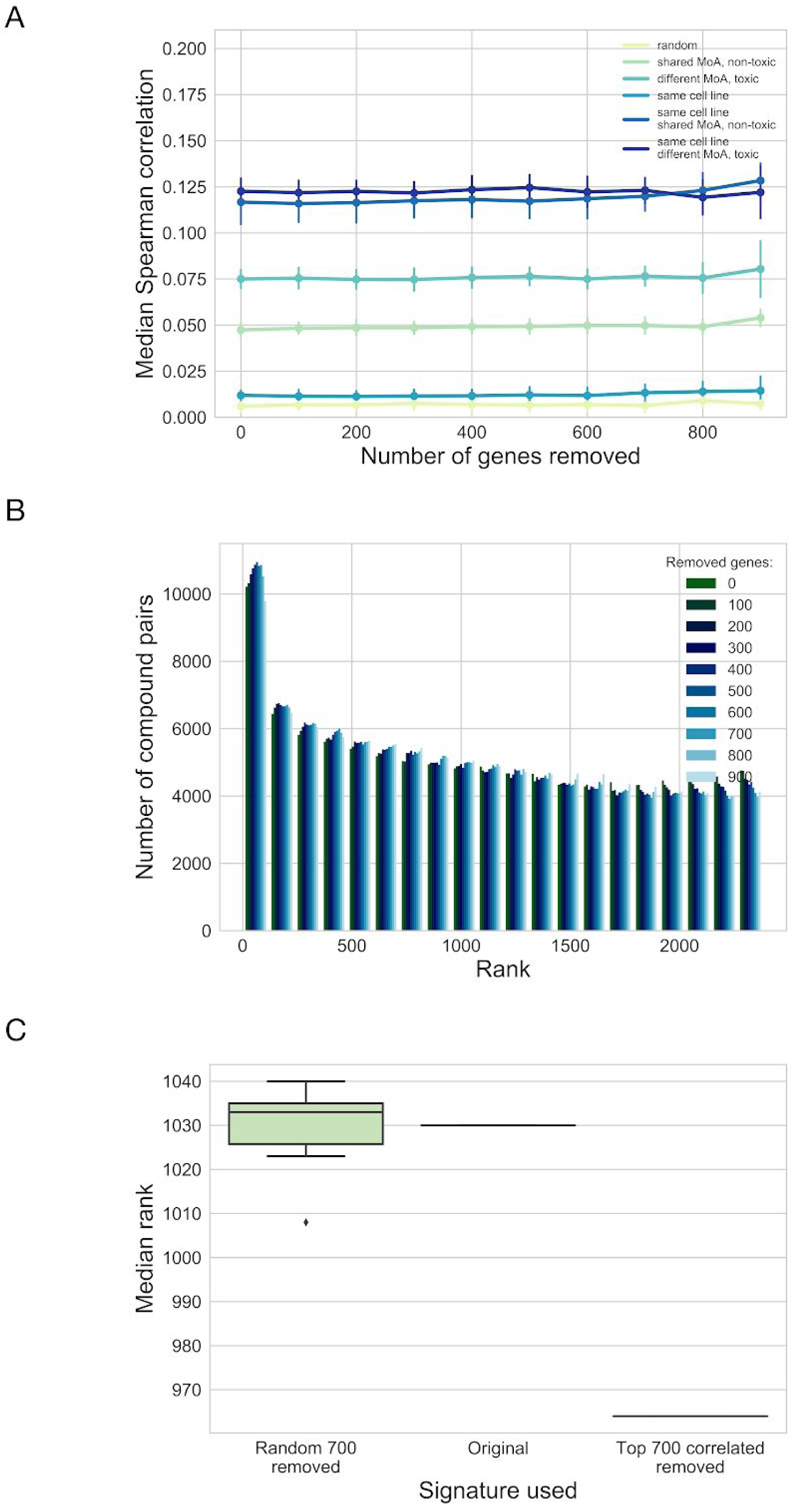
Effects of removing genes from L1000 signatures. (A) Effect of removing random genes on signature similarity. Random pairs of samples were taken from the CTRP-L1000–24h signatures with the following constraints (color code): absolutely random, non-toxic (cell viability>0.8) compounds with shared MoA and toxic (cell viability<0.6) compounds with different MoA. Signatures were sampled cell line irrespective or from the same cell line. Random n (x axis) genes were removed from the signatures before similarity calculation (1,000 sample pairs repeated 10 times, mean +/- SD). (B) Effect of removing cell death correlated genes on MoA discovery. Average signatures for 2485 compounds from LINCS-L1000-MoA dataset was calculated using the MODZ method. For each compound, other compounds were ranked based on signature similarity (Spearman correlation). Histogram (full distribution) shows the ranks of compounds with shared MoA. Top n (color codes) genes with highest absolute Pearson correlation with cell viability were removed before average signature and signature similarity calculation. (C) Effect of removing random genes on MoA discovery. Average signatures for 2485 compounds from LINCS-L1000-MoA dataset was calculated using the MODZ method. For each compound, other compounds were ranked based on signature similarity (Spearman correlation). Either 700 random genes (left, repeated 10 times) were removed or 700 genes with highest absolute correlation with cell viability were removed (right) or original (full) signatures were used for average signature and signature similarity calculations. Median ranks are shown for each case.

**Supplementary Figure 4 -.**
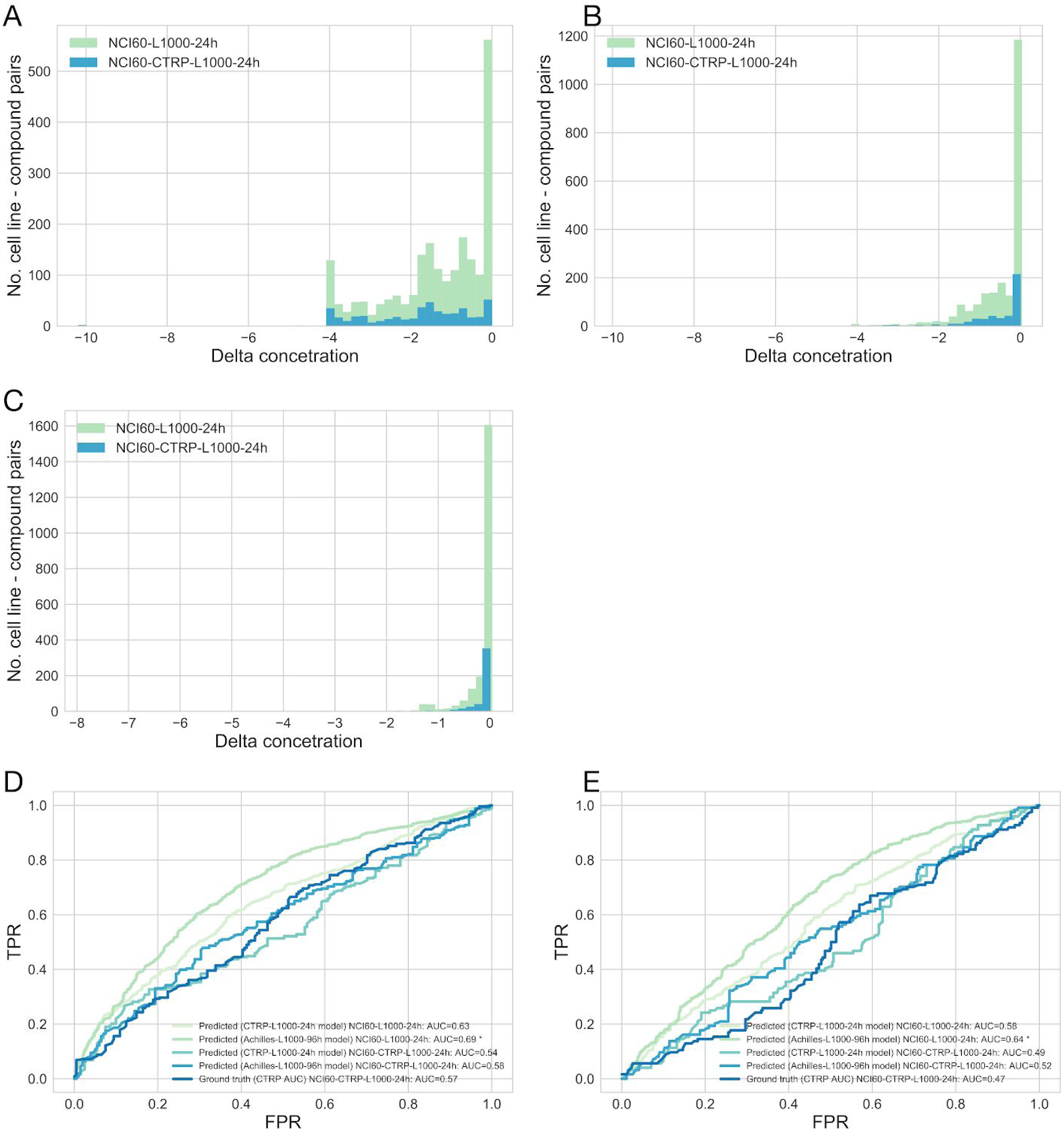
Distribution of toxic compounds and additional validation in the NCI60 dataset. (A-C) Distribution of effective and ineffective drugs in the NCI60-L1000–24h dataset. Delta concentration was defined as NCI60 sensitivity metric (GI50, TGI or LC50) - maximal used concentration. Delta concentration 0 means ineffective drug on the investigated cell line (as the drug does not have effect in the used concentration range). Used cell viability metrics are GI50 (A), TGI (B) and LC50 (C). (D,E) ROC analysis of the prediction performance of linear models on NCI60 data. Cell viability was predicted for the intersection of NCI60 and LINCS-L1000 or for the intersection of NCI60, CTRP and LINCS-L1000 datasets (NCI60-L1000–24h and NCI60-CTRP-L1000–24h respectively) using linear models trained on CTRP-L1000–24h or Achilles-L1000–96h data. Either these predicted cell viability values or the known AUC values from CTRP screen (CTRP AUC) were used to predict the binarised (effective / ineffective in the investigated concentration range) TGI (D) or LC50 (E) values from NCI60 (*: p<1e-5 for difference between AUCs for CTRP-L1000–24h and Achilles-L1000–96h).

**Supplementary Figure 5 -.**
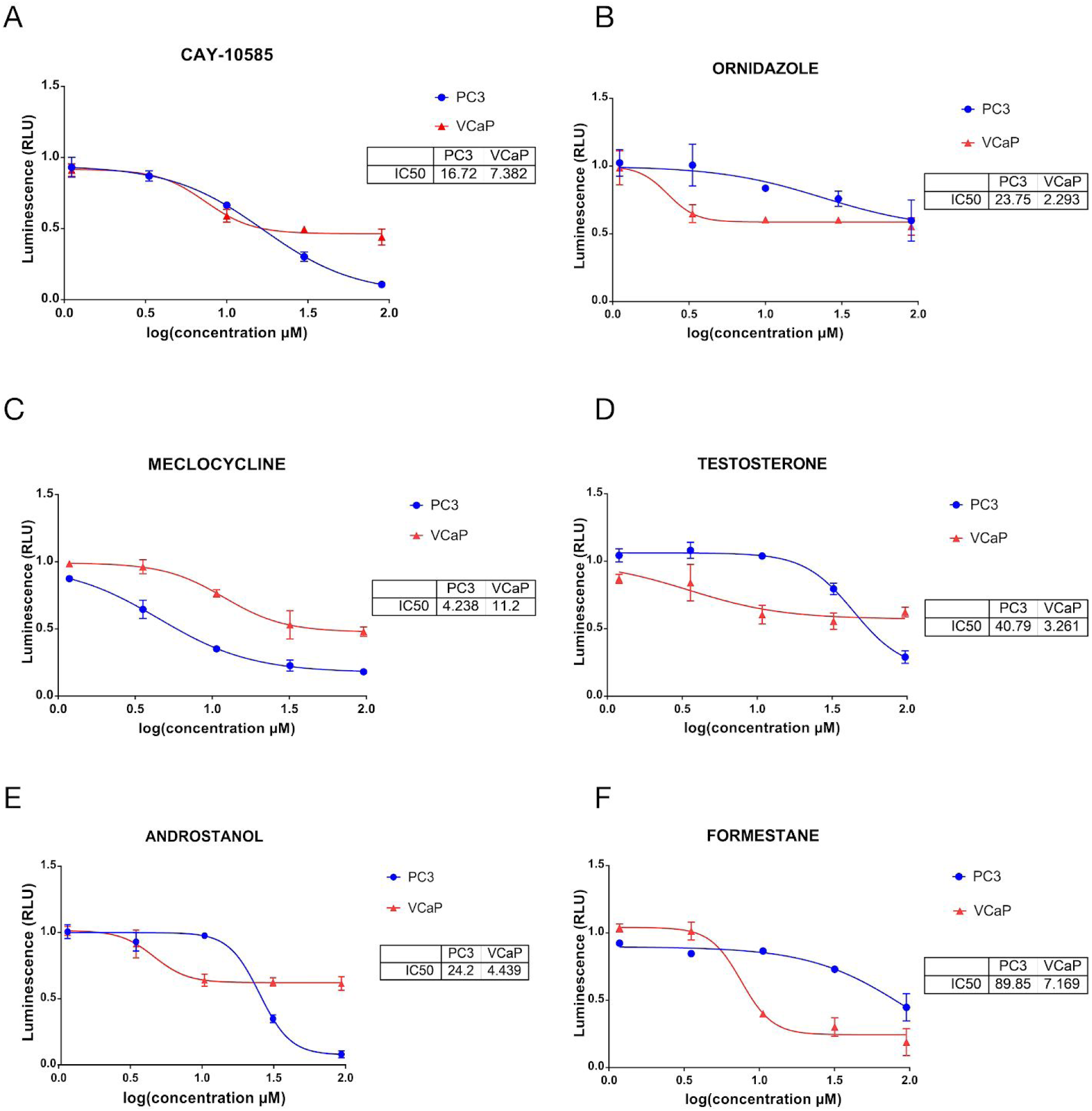
Experimental validation of prostate cancer cell line specific compound toxicity. (A-F) Dose response curves for experimentally tested compounds in PC3 and VCaP cell lines. Cell viability was measured in triplicates, after 48 hours incubation with tested compounds. Calculated IC50 values (GraphPad Prism) are shown in the inserts.

**Supplementary Table 1.** Descriptive statistics of the used datasets. Table includes data type (perturbation signature, cell viability or matched), number of data points, number of cell lines, number of compounds (small molecules or biologicals), shRNAs and time points (elapsed time between perturbation and measurement) for each used dataset.

|  | **Data type** | **Data points** | **Cell lines** | **Compounds** | **shRNAs** | **Time points** |
| --- | --- | --- | --- | --- | --- | --- |
| **LINCS-L1000** | signature | 591697 | 98 | 21921 | 18493 | 12 |
| **CTRP** | cell viability | 6171005 | 887 | 545 | 0 | 1 |
| **Achilles** | cell viability | 42893983 | 501 | 0 | 108718 | 1 |
| **CTRP-L1000** | matched | 16390 | 48 | 326 | 0 | 4 |
| **Achilles-L1000** | matched | 77230 | 11 | 0 | 12925 | 8 |
| **CTRP-L1000-3h** | matched | 1071 | 5 | 43 | 0 | 1 |
| **CTRP-L1000-6h** | matched | 7367 | 46 | 273 | 0 | 1 |
| **CTRP-L1000-24h** | matched | 7947 | 18 | 323 | 0 | 1 |
| **Achilles-L1000-96h** | matched | 57639 | 10 | 0 | 10733 | 1 |
| **Achilles-L1000-120h** | matched | 11431 | 2 | 0 | 11366 | 1 |
| **Achilles-L1000-144h** | matched | 7773 | 3 | 0 | 4180 | 1 |
| **LINCS-L1000-MoA** | signature | 103770 | 42 | 2485 | 0 | 1 |
| **LINCS-L1000-Chem** | signature | 320694 | 83 | 21921 | 0 | 8 |
| **NCI60** | cell viability | 3016553 | 159 | 52578 | 0 | 1 |
| **NCI60-L1000-24h** | matched | 2160 | 6 | 583 | 0 | 1 |
| **NCI60-CTRP-L1000-24h** | matched | 466 | 6 | 99 | 0 | 1 |
| **GDSC-L1000-24h** | signature | 21011 | 41 | 170 | 0 | 1 |

